# PRDX6 inhibits neurogenesis of neural precursor cells through downregulation of wdfy1 mediated TLR4 signal

**DOI:** 10.1101/086371

**Authors:** Mi Hee Park, Dong Ju Son, Kyoung Tak Nam, So Young Kim, Sang Yeon Oh, Min Ji Song, Hyung Ok Chun, Tae Hyung Lee, Jin Tae Hong

**Author notes:** Please address all correspondence to: Prof. Jin Tae Hong College of Pharmacy and Medical Research Center, Chungbuk National University, 194-31, Osongsaengmyeong 1-ro, Osong-eup, Cheongwon-gun, Chungbuk, Republic of Korea, 361-951.

## Abstract

Impaired neurogenesis has been associated with brain disorders. The role of peroxiredoxin 6 (PRDX6) in the neurodegenerative diseases is very controversial. To demonstrate the role of PRDX6 in neurogenesis, we compared neurogenesis ability and studied the molecular mechanisms. It was found that the neurogenesis of neural stem cells and expression of the marker protein were lowered in PRDX6 Tg-mice compared with non-tg mice. Moreover, the expression of wdfy1 was dramatically decreased in PRDX6-Tg mice, also, we observed that wdfy1 siRNA decreases the differentiation ability of primary neural stem cells to astrocyte and neuronal cells as well as PC12 cells. However, knockdown of PRDX6 recovered neurogenesis in the brain of PRDX6-Tg mice as well as PC-12 cells. We also showed that TLR4 was dramatically reduced in PRDX6 Tg mice as well as PC-12 cells and PRDX6 overexpression reduced neurogenesis was rescued after treatment of TLR4 siRNA. We further found that reduced TLR4 expression and neurogenesis was reversed in the neuron from PRDX6-Tg mice as well as PC12 cells by introduction of wdfy1 plasmid. Moreover, TLR4 siRNA reduced neurogenesis and wdfy1 expression. This study indicated that PRDX6 inhibits neurogenesis of neural precursor cells through TLR4 dependent downregulation of wdfy1.

## Summary statement

To demonstrate the role of PRDX6 in neurogenesis, we studied neurogenesis ability and molecular mechanisms. Our result showed that PRDX6 inhibits neurogenesis of neural precursor cells through TLR4 dependent downregulation of wdfy1.

## Introduction

Neurogenesis is impaired in neurodegenerative diseases, such as Alzheimer’s disease (AD) and parkinson’s disease (PD) (Curtis et al., 2007; Faure et al., 2011; Hoglinger et al., 2004). Extensive evidences have demonstrated that promotion of neurogenesis improves the symptoms of the diseases (Becker et al., 2007; Tchantchou et al., 2007). It has been demonstrated that anti-amyloid immunotherapy against the β-amyloid (Aβ) promotes recovery from AD and neurotoxicity by restoring neuronal population and cognitive functions in patients (Becker et al., 2007). Tchantchou et al also demonstrated that administration of EGb 761 recovered memory function by enhanced neurogenesis in the hippocampus of Tg2576 AD mouse model (Tchantchou et al., 2007). It is also reported that lithium improves hippocampal neurogenesis, thus recovers cognitive functions in APP mutant mice (Fiorentini et al., 2010). However, deficits of neural stem cell differentiation are associated with the development of neurodegenerative diseases. Familial alzheimer’s disease (FAD)-linked mutations of amyloid precursor protein (APP) transgenic mice showed impaired neurogenesis. Presenilin (PS)1 FAD mutant transgenic mice also showed in impaired neural precursor cell proliferation and survival (Elder et al., 2010; Winner et al., 2011). The double transgenic mice that expresses APP(KM670/671NL)/PS1(Δexon9) and mutant APPswe/PS1DeltaE9 in neurons showed neuritic dystrophy (Readhead et al., 2016) and impairments in neurogenesis (Demars et al., 2010). It has been also reported that PD-associated Leucine-rich repeat kinase 2 (LRRK2) mutation inhibits neuronal differentiation of neural stem cells (Bahnassawy et al., 2013). PTEN-induced putative kinase 1 (PINK1) deficiency mice also decrease brain development and neural stem cell differentiation (Choi et al., 2016). Recent study also demonstrated that Synuclein Alpha (SNCA) gene triplication that is involved in PD impairs neuronal differentiation and maturation of PD patient-derived induced pluripotent stem (iPS) cells (Oliveira et al., 2015). Moreover, recent study demonstrated that proliferating cells do not become mature neurons in the AD brain (Richetin et al., 2015), and the number of proliferating cells was reduced in brain of PD patient compared with controls (Regensburger et al., 2014). These results suggested that neurodegenerative diseases including AD and PD are associated with impaired neurogenesis.

Because of the multiple damage-response pathways in the central nervous system (CNS) regulated by the redox state, oxidative stress plays a major role in neuronal death caused by severe vulnerability of the brain to a lost oxidation-reduction balance (Nixon and Crews, 2002). It has been demonstrated that cell differentiation show marked correlations with cellular oxygen species levels in the nervous system. Oxidative stress is a deleterious condition leading to cellular death and plays a key role in the development and pathology of neurodegenerative diseases through impaired neurogenesis. In the pathophysiology of neurodegenerative diseases, such as PD, AD (Grote and Hannan, 2007), and also in other pathologies, such as ischemic stroke, a state of oxidative stress accompanied by the alteration of neurogenesis has been implicated (Wiltrout et al., 2007). Antioxidant reduces the deleterious activity of oxidative stress and potentially delays the development of neurodegenerative disease, the study of antioxidant enzyme in neurogenesis is very important for prevention of the development of neurodegenerative disease. The recent finding showed that when examined at 3-4 months of age, genetic deficiency of extracellular superoxide dismutase (SOD) was associated with a significant suppression of baseline neurogenesis and impaired hippocampal-dependent cognitive functions (Zou et al., 2012). Other study also demonstrated that lack of extracellular SOD in the microenvironment impacts radiation-induced changes in neurogenesis (Rola et al., 2007). It is also reported that antioxidant proteins, metallothionein recovered brain injury through increasing neurogenesis (Penkowa et al., 2006). Metallothionein null mice display impaired brain parenchyma recovery following injury, whereas transgenic metallothionein overexpression transgenic and exogenous administration of metallothionein lead to less damage through enhancement of neurogenesis (Giralt et al., 2002). A neuroprotective role of metallothionein has also been shown in other models of CNS injury, such as experimental autoimmune encephalomyelitis (Penkowa et al., 2001). Other study also demonstrated that malpar1 KO mice which displays enhanced oxidative stress in the hippocampus, showed abnormalities of hippocampal neurogenesis (García-Fernández et al., 2012). Catalase overexpressed mice improve neurogenesis, and has neuroprotective effect against radiation induced improvement of hippocampal neurogenesis (Liao et al., 2013). Peroxiredoxin (PRDX)s are also well studied in neural progenitor cell differentiation. Cytoplasmic PRDX1 and PRDX2 actively participate in the maintenance of embryonic stem cell stemness through opposing ROS/JNK activation during neurogenesis (Kim et al., 2014). PRDX4 ablation causes premature neuron differentiation and progenitor depletion (Yan et al., 2015). Besides of other PRDXs, the role of PRDX6 in the neurogenesis has not been reported yet. However, several our studies have demonstrated that PRDX6 overexpression even accelerates the development of AD, PD as well as EAE. Thus, we could speculate that PRDX6 may cause impaired neurogenesis in these accelerating processes.

Several signal pathways regulated by antioxidant enzymes have been reported for the control of neurogenesis. Nrf2-driven Heme oxygenase1 associated with (Wnt)/β-catenin signaling was causally related to the impairment of proliferation and differentiation of neural precursor cells (L’Episcopo et al., 2013). Other report suggested that excessive NO generation through activation of CREB pathway modulates neuronal differentiation and development (Okamoto and Lipton, 2015). Several studies suggested that these critical pathways are regulated by TRIF to Toll-like receptor (TLR) 4 pathway. Rebeca et al demonstrated that TLR4 activates the β-catenin pathway (Kamo et al., 2013). Moreover, TLR4 induces CREB signalling regulating early survival, neuronal gene expression and morphological development in adult subventricular zone neurogenesis (Herold et al., 2011). TLRs have recently been documented to be implicated in neurogenesis in mammalian (Rolls et al., 2007) and brain development (Ma et al., 2007). TLR2 and TLR4 are found on adult neural stem/progenitor cells and have distinct and opposing functions in neural stem cells proliferation and differentiation both in vitro and in vivo (Rolls et al., 2007). TLR2 deficiency in mice impaired hippocampal neurogenesis, whereas the absence of TLR4 resulted in enhanced proliferation and neuronal differentiation (Rolls et al., 2007). In vitro studies further indicated that TLR2 and TLR4 directly modulated self-renewal and the cell-fate decision of NPCs. These data indicate that TLR pathway, and TLR associated signal pathway could be significant in the control of oxidative stress-induced neurogenesis.

In the preliminary study, we identified that PRDX6 overexpression causes impairment of neurogenesis, and several factors were changed in the neural stem cells isolated from PRDX6 overexpressed transgenic mice. Among them, Wdfy1 is dramatically decreased in neural stem cells isolated from PRDX6 overexpressed transgenic mice compared with non-tg mice. Wdfy1 have identified WD-repeat- and FYVE-domain-containing protein 1 (wdfy1) as a downstream of the Neuropilin 2 (NRP2) axis (Dutta et al., 2016). Wdfy1 is involved in placental development and for the maintenance of hematopoietic stem cells (Bennett et al., 2015). Interestingly, wdfy1 has been shown to recruit signaling adaptor, TLR3 and TLR4 (Hu et al., 2015), and thereby potentiating signaling necessary for the neural stem cell differentiation. Although the possibility of Wdfy1 in development of neural stem cell is suggested, however, its role on PRDX6-induced neurogenesis and possible mechanisms are not clear. Therefore, we investigated a link between PRDX6 and Wdfy1/TLR signal for the control of neural stem cell differentiation and action mechanisms in PRDX6 induced impairment of neurogenesis using PRDX6 overexpressed transgenic mice.

## Results

### Effect of PRDX6 on the neurite outgrowth of neural stem cells

To study the role of PRDX6 on the spontaneous differentiation of neural stem cells, we cultured primary neural stem cells isolated from cortex of E15 days mouse embryos of non-tg or PRDX6 Tg-mice. The cultured cells formed neurospheres at 5 days in vitro. We showed that neural stem cells from PRDX6 Tg mice resisted spontaneous differentiation for a prolonged time. PRDX6 mice showed impaired differentiation, and were unsuccessful in forming a high-quality network of neural cells. We observed that neural stem cells from RPDX6 overexpressed mice had significantly shorter length of primary and secondary neurites as well as lower number of neurites per cell as compared to that of non-tg (Fig. 1A). To understand the contribution of PRDX6 on differentiation of neural stem cells into the specific cell types, we determined specific neuronal cell differentiation. We found that the differentiated neuron and glial cells (Fig. 1B) are much lower in PRDX6 Tg mice. We also showed that the expression of tubulin β-III (TUBBIII) that is a neuronal cell marker was decreased, and the expression of neurofilament (NF) that is a major component of the neuronal cytoskeleton was also decreased in PRDX6 overexpressed mice. Moreover, the expression of glial fibrillary acidic protein (GFAP), an astrocyte specific marker was also decreased in PRDX6 Tg mice. To check if the effect of PRDX6 was similar in non-stem cells, PC12 cells were differentiated for 5 days upon stimulation with NGF (100 ng/ml) after the introduction of PRDX6 plasmid. PC12 has been previously used as an instructive model for studying the underlying the neuronal differentiation in response to NGF (Gupta et al., 2011). We showed that neurite-outgrowth and branching of PC12 cells were stimulated by the treatment of NGF, and this effect was inhibited by the treatment of PRDX6 plasmid. In a quantified data, the average number of neurites per cell was much lower in PRDX6 plasmid treated cells as compared to control cells (Fig. S1A). Thus, PRDX6 could be involved in neuronal differentiation both in the PC12 cell line as well as neural stem cells. PRDX6 has dual enzyme activity, iPLA2 and glutathione peroxidase activity. Our recent study indicated that PRDX6 accelerate the AD development through iPLA2 activation (Yun et al., 2013). So, we defined whether iPLA2 is involved in impaired neurogenesis. We observed that differentiation ability from neural stem cells to neuronal cell and astrocyte was reversed after treatment of PRDX6 inhibitor (Fig.1C, left panel) and iPLA2 siRNA (Fig.1C, right panel), similar effects was found in the differentiation of PC12 cells (Fig. S1B). From these result, we suggested that the PRDX6 involved in the differentiation of specific cell types of brain.

**Fig.1.**
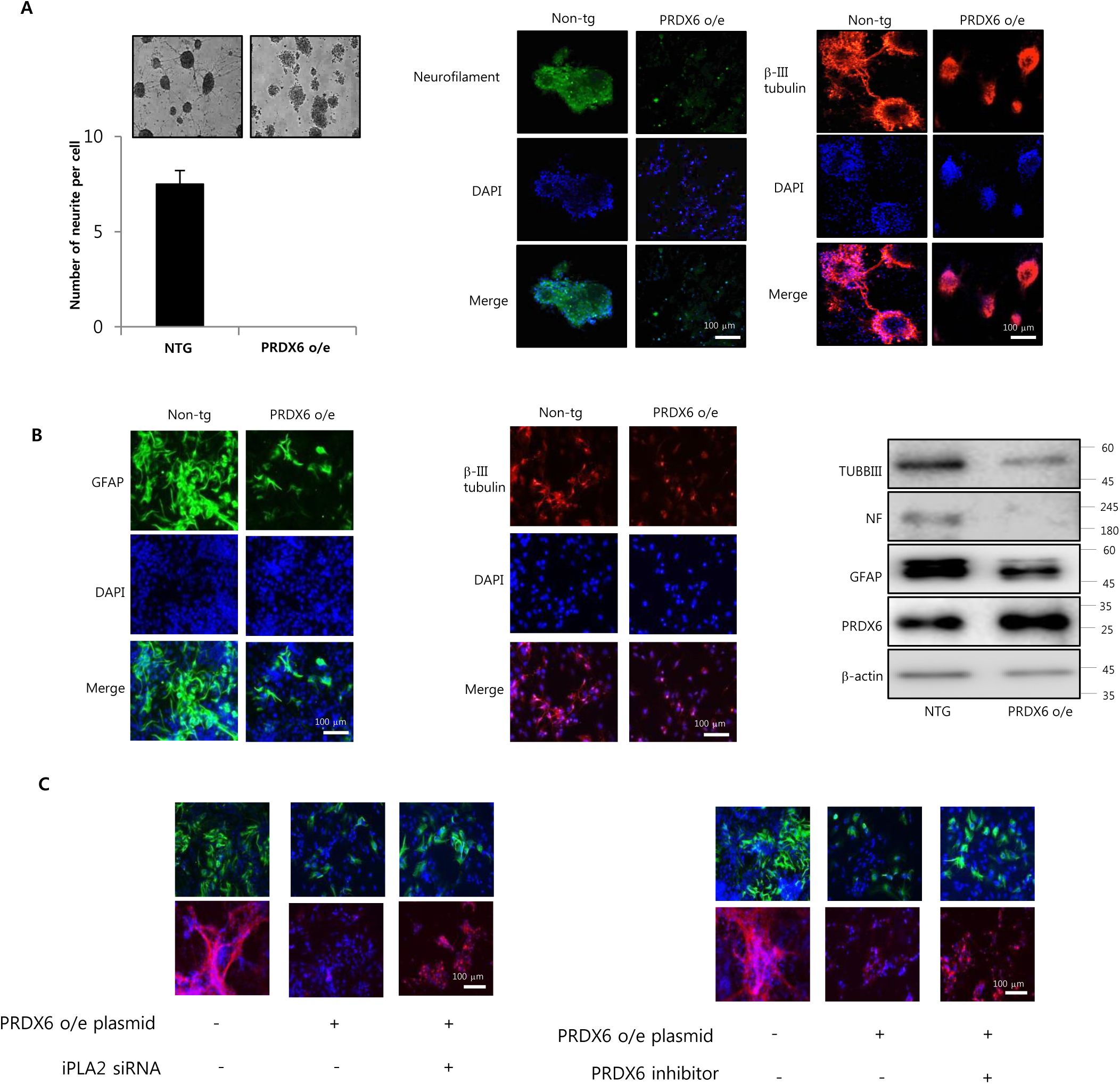
**Effect of PRDX6 on the differentiation of neural stem cells. A**, Neural stem cells were isolated from embryonic day 15 forebrain germinal zones from Non-tg or PRDX6-overexpressed tg mice. Bulk cultures were established in the complete medium, then after 24hr, the medium was changed with Neurobasal medium containing supplements. The cultured cells formed neurospheres at 5 days in vitro. **B**, Neural stem cells were isolated, then differentiated into astrocytes and neuronal cells in vitro as described in materials and methods. Western blot analysis confirmed the expression of beta III-tubulin, GFAP, NF and PRDX6 in terminally differentiated neurons and astrocytes. β-actin was internal control. **C,** Neural stem cells were transfected with PRDX6 for 24hr, and then transfected with iPLA2 siRNA (100 nM) or treated with PRDX6 inhibitor for 24hr, then differentiated into astrocytes and neuronal cells in vitro as described in materials and methods and then immunostained with GFAP or beta III-tubulin antibodies. Images shown are representative microscopy images (n=5 each).

### Involvement of wdfy1 in the differentiation of neural cell lineage

To determine what factors are involved in PRDX6 related neural stem cell impairment, we did microarray experiment. In microarray data, several factors are changed. Among them, wdfy1 is dramatically changed in PRDX6 tg mice derived neural stem cells (Fig. 2A). Eventhough other factors such as Gm2964, Vmn1r80, Clca1, Pisd-ps3 and Gm10002 were greatly changed, siRNA study of these factors did not affect on neurogenesis (data not shown). Also, we confirmed that wdfy1 expression was dramatically decreased in PRDX6 tg mice by using RT-PCR (Fig. 2B), Western blotting (Fig. 2C) and immunohistochemistry (Fig. 2D). Few studies have been reported on the role of WDFY1 in neurogenesis. First, we found that wdfy1 expression was increased in PC12 cells after treatment of NGF and neural stem cells after differentiation (5 days) (Fig.S2A). We showed that neurite-outgrowth and branching of PC12 cells were stimulated by treatment of NGF, and this effect was inhibited by treatment of wdfy1 siRNA. Average number of neurites per cell was much lower in wdfy1 siRNA treated cells as compared to control cells in a quantified data (Fig. 3A) accompanied with reduced expression of GFAP, TUBBIII and TH (Fig.S2B, left panel). We also observed that wdfy1 siRNA decreased differentiation ability of neural stem cells to astrocyte and neuronal cells was also decreased (Fig. 3B). Next, we performed the experiment to show the differentiation ability of wdfy1 in *in vivo*. We injected the neural stem cells transfected by wdfy1 siRNA in SVZ region. In immunohistochemical staining, differentiated neuronal cells and astrocytes were dramatically decreased in wdfy1 siRNA (Fig. 3C) accompanied with reduced expression of GFAP, TUBBIII and TH (Fig.S2B, right panel). We also found that the impaired differentiated astrocyte (Fig.4A) and neuronal cells (Fig.4B) from neural stem cell induced by PRDX6 overexpression was rescued by introduction of wdfy1 plasmid by immunofluorescence staining and Western blotting.

**Fig.2.**
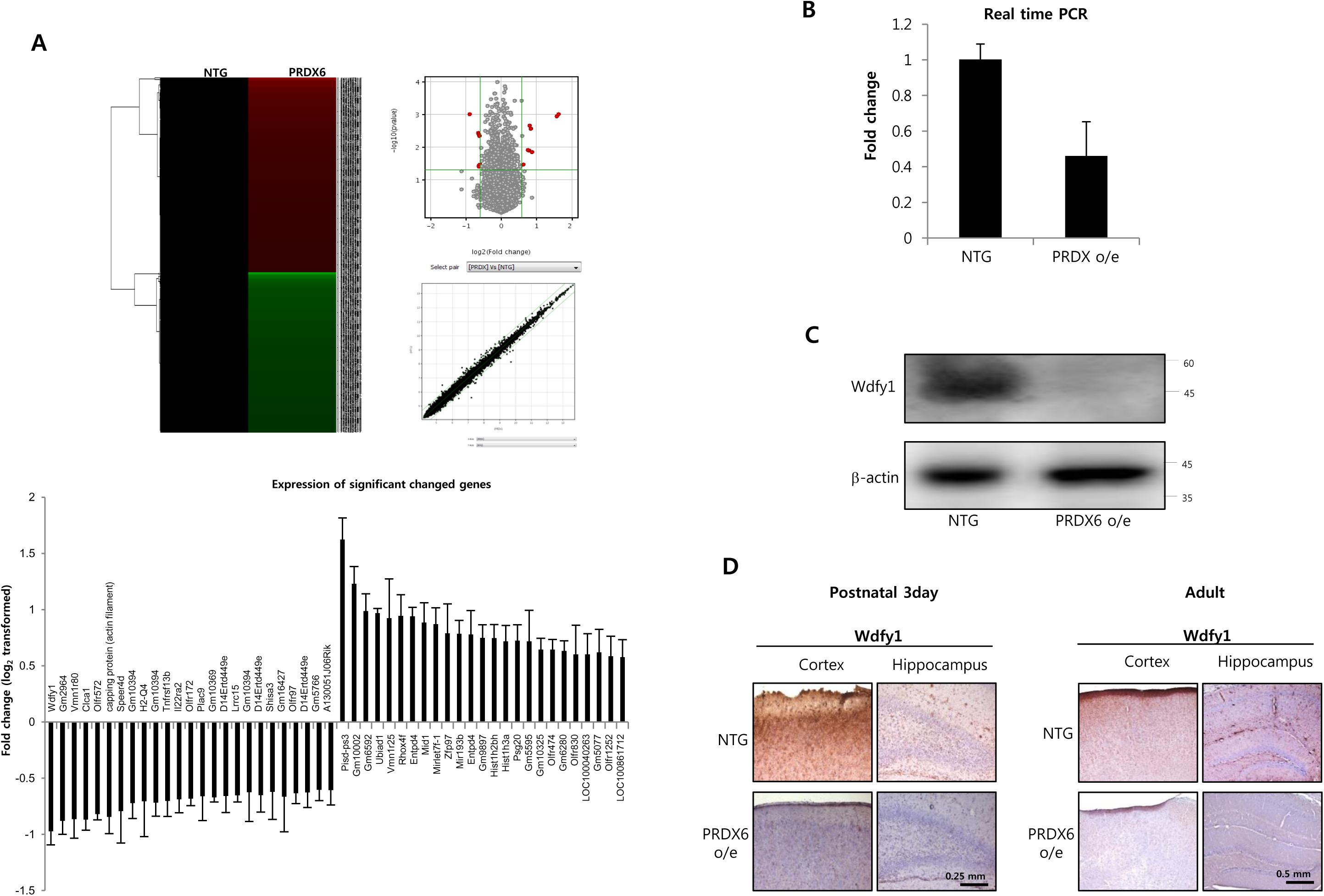
**A, Global gene expression profiles in the neural stem cells isolated from PRDX6 o/e Tg mice.** Total RNA were obtained from the neural stem cells after 5days of differentiation. Affymetrix GeneChip Mouse Gene 2.0 ST arrays containing 770, 317 mouse gene probes were used. Scatter plots show normalized intensities of each probe (**A**). The expression of wdfy1 mRNA was determined by qPCR (**B**) and western blotting (**C**). **D**, Paraffin sections of brain from PRDX6 o/e Tg or non-tg mice were stained with antibody against PRDX6. Images shown are representative microscopy images (n=5 each).

**Fig.3.**
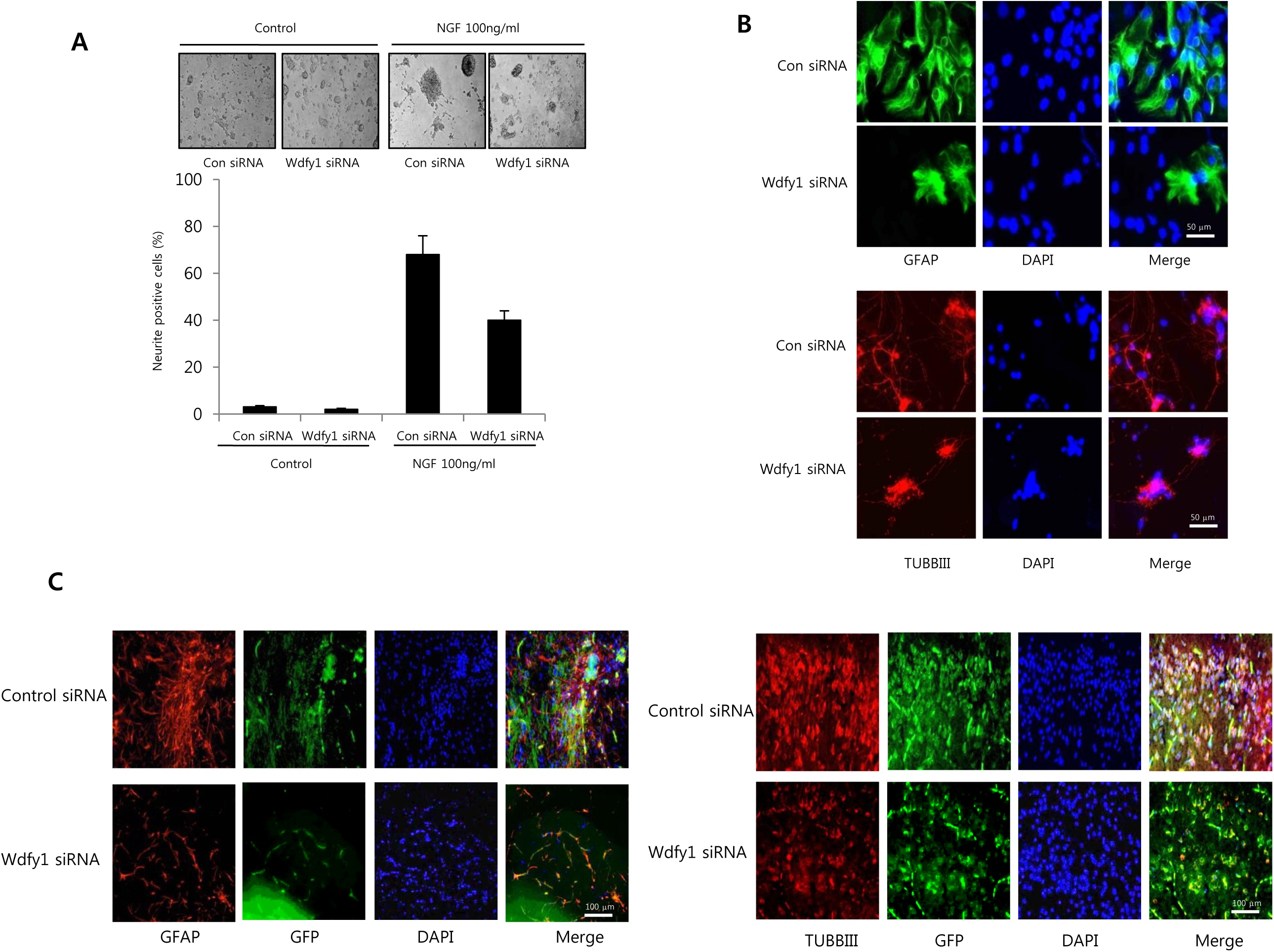
**Effect of knockdown of wdfy1 on the differentiation of neural stem cells. A**, PC12 cells were differentiated for 5 days upon stimulation with NGF (100 ng/ml) after introduction of wdfy1 siRNA (100 nM). To study neurite outgrowth, the medium was changed to RPMI containing 100 ng/ml NGF. The cells were further cultured for 5 days. Cells with at least one neurite longer than two-body length were counted as neurite positive. At least 500 cells were counted for each group performed in triplicate. **B**, Neural stem cells were transfected with wdfy1 siRNA for 48hr, then differentiated into astrocytes and neuronal cells in vitro as described in materials and methods and then immunostained with GFAP or beta III-tubulin antibodies. **C**, Neural stem cells tranfected with wdfy1 siRNA, and then injected in SVZ region of ICR mice. After 2 weeks, differentiated astrocyte or neuronal cells were stained with GFAP or beta III-tubulin. Images shown are representative microscopy images (n=5 each).

**Fig.4.**
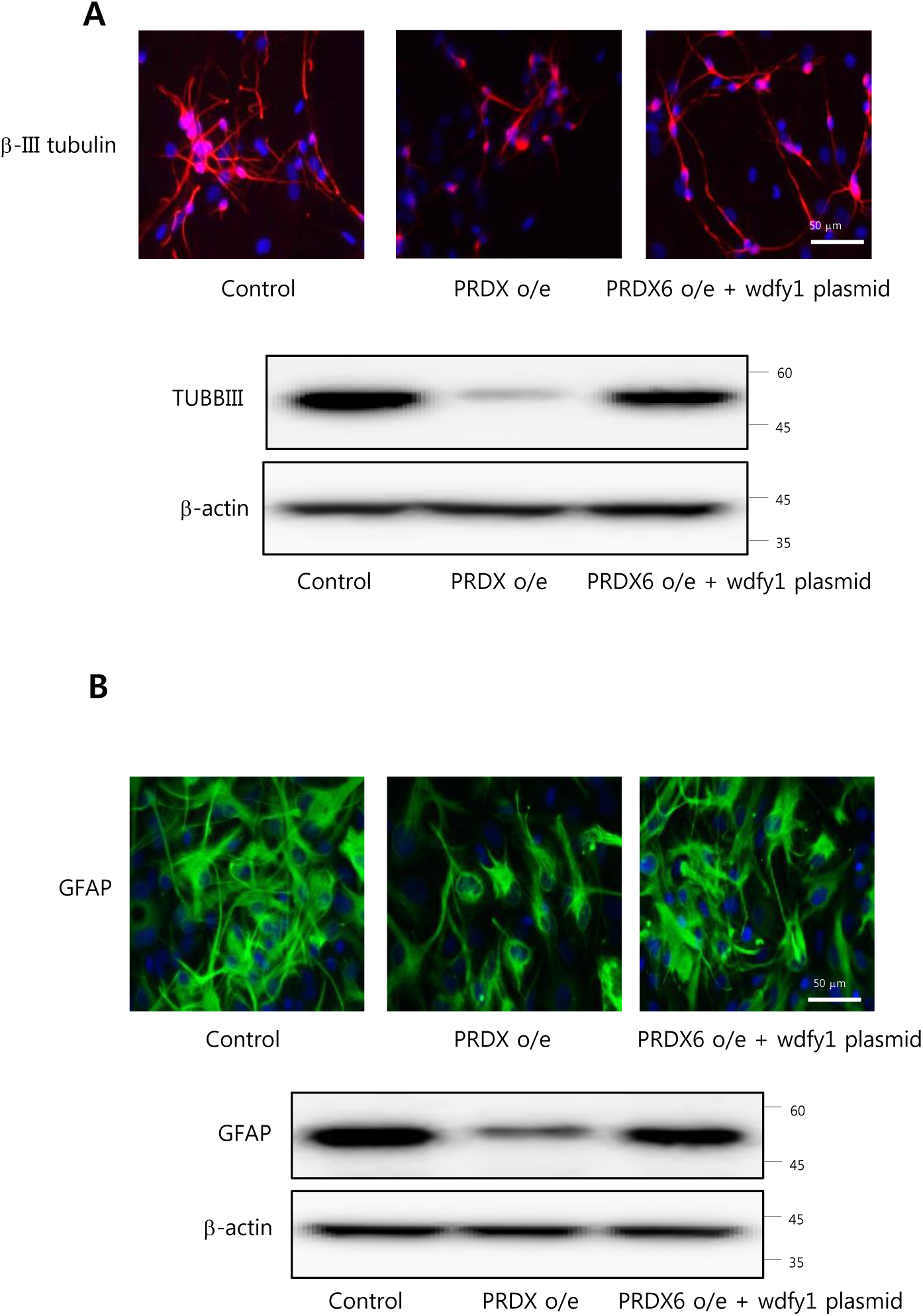
**Effect of wdfy1 on the PRDX6-induced impaired differentiation of neural stem cells.** Neural stem cells isolated from non-tg or PRDX6 o/e tg mice were transfected with wdfy1 plasmid. Then neural stem cells were differentiated into neuronal cells (**A**) and astrocytes (**B**) in vitro as described in materials and methods and then immunostained with beta III-tubulin or GFAP antibodies. β-actin was internal control. Images shown are representative microscopy images (n=5 each).

### Involvement of TLR pathways in the wdfy1 dependent impairment of neuronal cell differentiation in PRDX6 overexpressed-Tg mice

To determine the involvement of TLR pathways are involved in wdfy1 dependent PRDX6 induced impaired neurogenesis. Among several TLR genes, expression of TLR4 and TLR5 was significantly decreased in neural stem cells from PRDX6 Tg mice compared with non-tg mice (Fig.5A), however, expression of other TLRs were not significantly changed in PRDX6 Tg mice (Fig.S3). Next, we confirmed the expression patteren in these mice by Western blotting and immunohistochemistry. We showed that the TLR4 was dramatically reduced in PRDX6 Tg mice (Fig.5B). We also determined the expression pattern of TLR4 depends on the development of brain. Brains were isolated at embryonic 15 days, embryonic 17 days, postnatal 0 day, postnatal 3 days and postnatal 5 days, and then Western blotting. We showed that the TLR4 expression was increased depends on the development of brain accompanied with increased wdfy1, but decreased PRDX6 (Fig.6A). In addition, we confirmed the expression of other PRDXs subtypes. PRDX1 was little decreased, PRDX2 and PRDX4 were increased, PRDX3 was not detected, and PRDX5 was not changed, but PRDX6 was dramatically decreased in a postnatal day-dependent manner by RT-PCR (Fig.6B). We also observed that TLR4 was dramatically increased in a postnatal day-dependent manner by immunohistochemistry (Fig.6C). Next, we showed the effect of knockdown of TLR4 on the differentiation of neural stem cells. We showed that differentiated astrocyte and neuronal cells was reduced by introduction of TLR4 siRNA demonstrated by IF staining and Western blotting (Fig.6D). We further found that reduced TLR4 expression was reversed by introduction of wdfy1 plasmid (Fig.6E). From these results we suggest that TLR4 is critical for the development of neural stem cells and this pathway is involved in the wdfy1 dependent neurogenesis in PRDX6 overexpressed mice.

**Fig.5.**
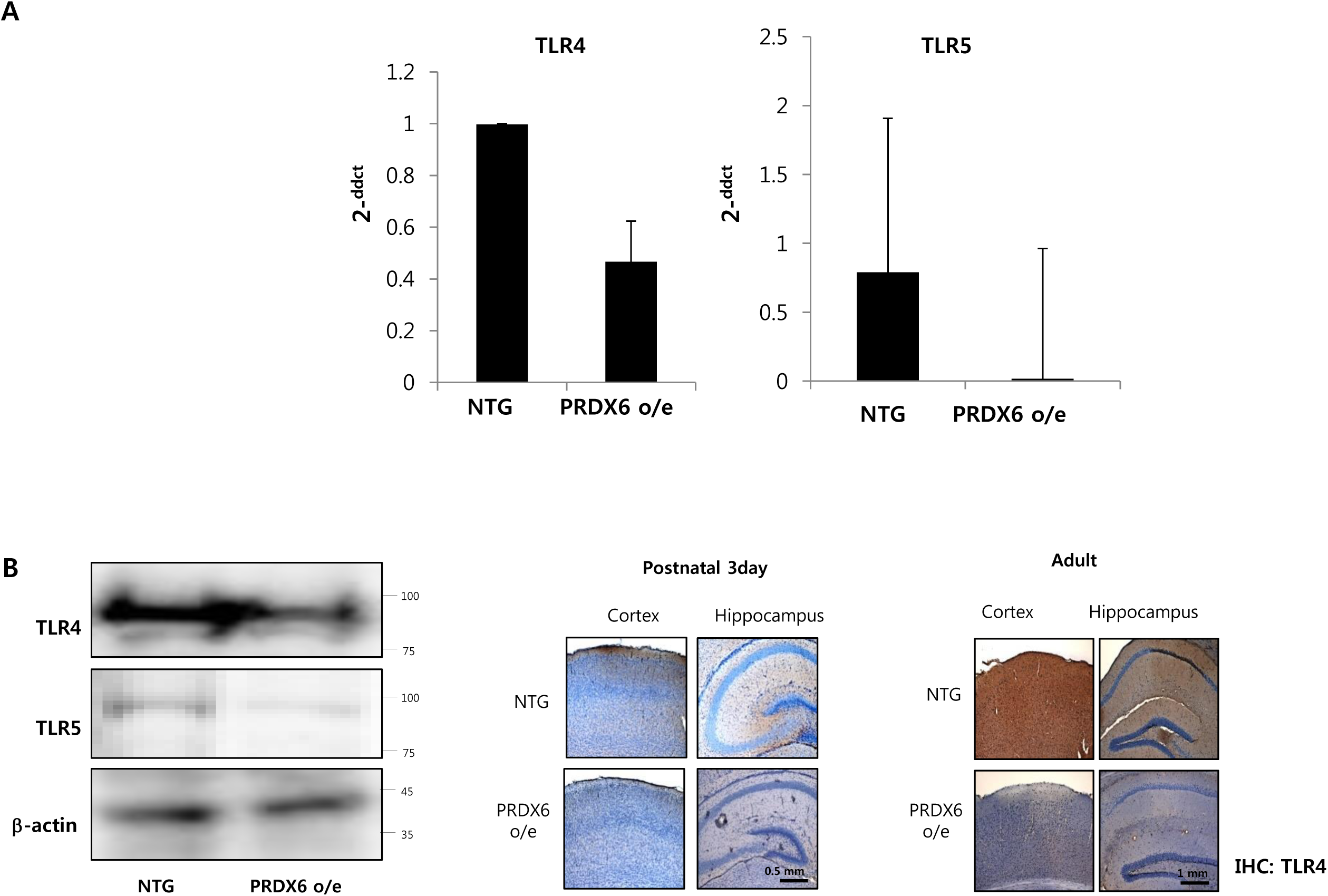
**Effect of PRDX6 on the expression TLR genes.** Total RNA were obtained from the neural stem cells after 5 days of differentiation. Then, the expression of TLR4 and TLR5 in neural stem cells was measured by qPCR (**A**) and Western blotting (**B**), and TLR4 by immunohistochemistry (**C**). Images shown are representative microscopy images (n=5 each).

**Fig.6.**
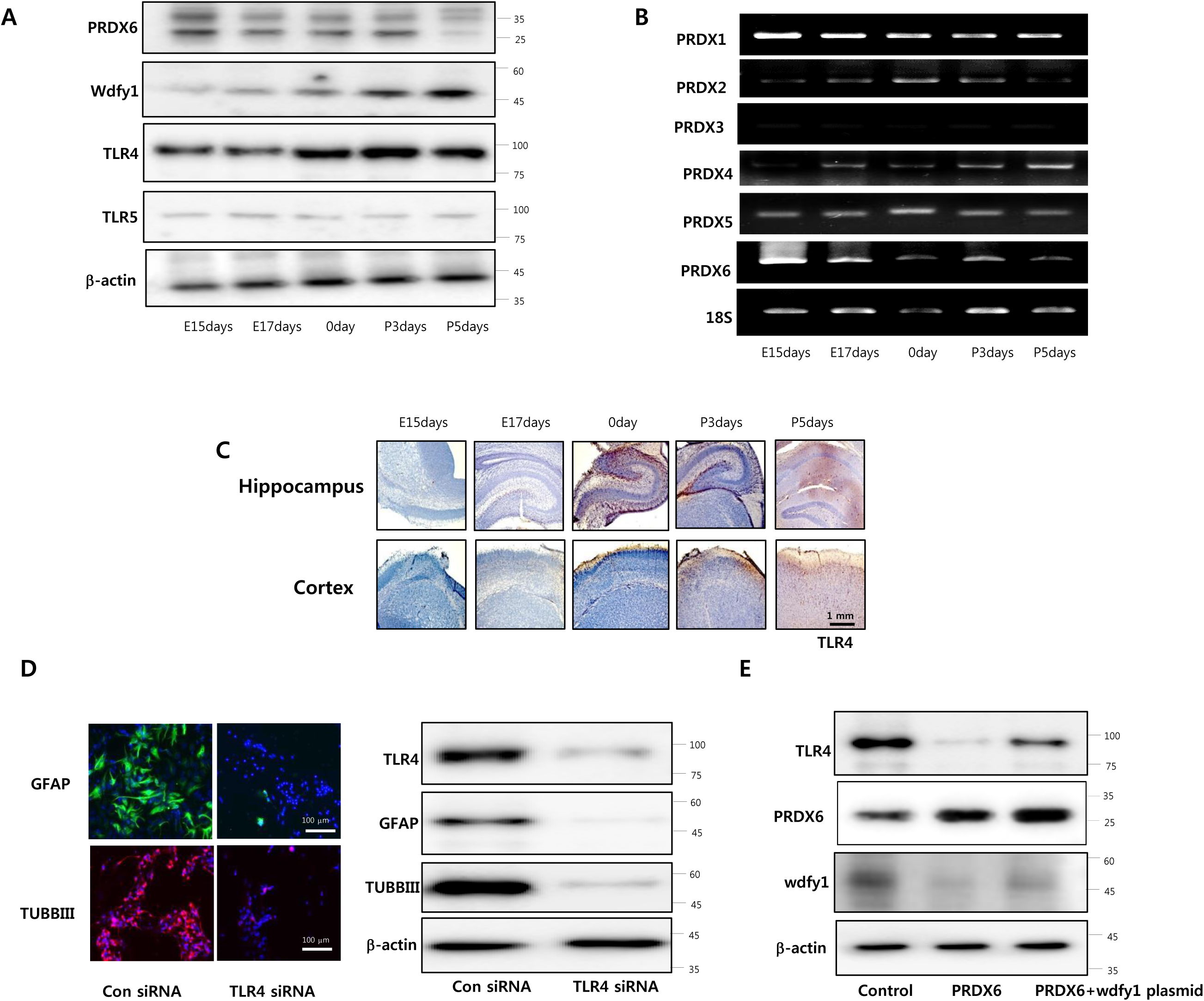
**Expression pattern of TLR4 and wdfy1 on the development of brain of PRDX6 overexpressed mice.** Brains were isolated at embryonic 15 days, embryonic 17 days, postnatal 0 day, postnatal 3 days and postnatal 5 days, and then western blotting was performed with PRDX6, wdfy1, TLR4 and TLR5 antibodies (**A**), and RT-PCR was performed with PRDX1~6 specific primers (**B**). **C**, Paraffin sections of brains were stained with antibody against TLR4 antibody. **D**, Effect of knockdown of TLR4 on the differentiation of neural stem cells. Neural stem cells were transfected with TLR4 siRNA. Then neural stem cells were differentiated into neuronal cells and astrocytes in vitro as described in materials and methods and then immunostained with GFAP or TUBBIII antibodies or western blotted with TLR4, GFAP or TUBBIII antibodies. **E**, Neural stem cells isolated from non-tg or PRDX6 o/e tg mice were transfected with wdfy1 plasmid. Then neural stem cells were differentiated and then western blotted with TLR4, PRDX6 or wdfy1 antibodies. β-actin was internal control. Images shown are representative microscopy images (n=5 each).

## Discussion

We demonstrated here that the role of PRDX6 on the impaired neurogenesis through downregulation of Wdfy1 mediated TLR4 pathway by using PRDX6 Tg mice derived neural stem cells and knockdown of PRDX6 in cultured cells. Recent our studies demonstrated that PRDX6 accelerate the development of AD through increased amyloidogenesis (Yun et al., 2013). We also recently found that PRDX6 Tg mice had a much higher loss of dopaminergic neurons resulting promote the development of PD (Yun et al., 2015). Other report also suggested that alterations in PRDX6 levels are associated with neurodegenerative disease such as Pick disease and sporadic Creutzfeldt-Jacob disease (Gao et al., 2005). It has been suggested that neurogenesis is impaired in these neurodegenerative diseases. Oxidative stress plays a major role in neuronal death caused by severe vulnerability of the brain to a lost oxidation-reduction balance (Nixon and Crews, 2002). In the pathophysiology of neurodegenerative diseases, a state of oxidative stress influences on neurogenesis (Wiltrout et al., 2007). It has been demonstrated that lack of extracellular SOD in the microenvironment impacts radiation-induced changes in neurogenesis (Rola et al., 2007). Overexpression of other antioxidant enzymes such as catalase and glutathione peroxidase also increases neurogenesis, and thus preventing neuronal injury. PRDX1, PRDX2 and PRDX4 overexpression is also related with neurogenesis (Kim et al., 2014; Yan et al., 2015). Cytoplasmic PRDX1 and PRDX2 actively participate in the maintenance of embryonic stem cell stemness through opposing ROS/JNK activation during neurogenesis (Kim et al., 2014). PRDX4 ablation causes premature neuron differentiation and progenitor depletion (Yan et al., 2015). Besides of other PRDXs, the role of PRDX6 in the neurogenesis has not been reported yet. Although the impairment of neurogenesis by oxidative stress is associated with the accelerative effects on the neurodegenerative diseases, the role of PRDX6 on neurogenesis, and its mechanisms are not well studied.

In our study, PRDX6 overexpressed mice showed impairment of neurogenesis. In addition, overexpressed PRDX6 increased PC12 cell differentiation, moreover knockdown of PRDX6 recovered neurogenesis. We also confirmed that the expression of PRDX6 was gradually decreased in the embryo development. However, the inhibitory effects on neurogenesis did not recovered by the treatment of antioxidants such as Vit E and glutathione (data not shown). We also found that PRDX6 overexpressed mice did not inhibit neurogenesis after iPLA2 knockdown or treatment of PRDX6 inhibitor. In this regard, it is noteworthy that overexpression of PRDX6 accelerated neurodegenerative diseases by the increase of iPLA2 activity covering antioxidant capability in the brain. Moreover, although the impairment of neurogenesis by oxidative stress is associated with the accelerative effects on the neurodegenerative diseases, the role of PRDX6 on neurogenesis, and its mechanisms are not well studied. In this study, we checked all subtypes of PRDXs at the developmental stages. PRDX1 was little decreased, PRDX2 and PRDX4 were increased, PRDX3 was not detected, and PRDX5 was not changed, but PRDX6 was dramatically decreased in a postnatal day-dependent manner by RT-PCR. These data indicate that the impairment of neurogenesis by iPLA2 activity through PRDX6 overexpression could be associated with acceleration of the development of neurodegenerative diseases.

To determine what factors are involved in PRDX6 related neural stem cell impairment, we did microarray experiment. In microarray data, we identified that several factors were changed in differentiated neural stem cells isolated from PRDX6 overexpressed Tg mice compared with non-tg mice. Among them, we showed that Wdfy1 was dramatically decreased in neural stem cells isolated from PRDX6 overexpressed Tg mice. However, other factors did not change neurogenesis after treatment of siRNA of these factors. We thus confirmed the reduced expression of wdfy1 in neural stem cells of PRDX6 mice by qPCR, Western blotting and immunohistochemistry analysis. The role of Wdfy1 was not fully understood, because few studies have been reported regarding the role of WDFY1 on neurogenesis. WDFY1 functions as downstream of NRP2 which has important role for gangliogenesis, axon guidance and innervation in sympathetic nervous system (Maden et al., 2012). Other study also demonstrated that loss of wdfy3 in mice alters cerebral cortical neurogenesis, suggesting that wdfy3 is essential to cerebral expansion and functional organization (Orosco et al., 2014). Other reports have indicated a role of WDFY1 in maintaining hematopoietic stem cells. It was shown that over-expression of WDFY1 abrogated the early endosomal maturation, and thereby hindered the fusion of autophagosomes with the late endosomes for the formation of autolysosomes (Dutta et al., 2016). In the present study, we found that decreased neurogenesis in PRDX6 overexpressed mice was recovered by wdfy1 expression, however, wdfy1 knockdown inhibited NGF-induced PC12 cell differentiation, and normal neurogenesis. To determine the role of wdfy1 on neurogenesis, we stereotaxically injected the wdfy1 shRNA transfected-neural stem cells into the dorsal horn of the SVZ of 10-wk-old ICR mice. After 2 weeks following the injection, the injected neural stem cells were differentiated to astrocyte and neuron cells in the SVZ region of the brain by immunofluorescence staining. We showed that GFAP positive cells that are astrocyte cell markers are decreased in the wdfy1 shRNA transfected cell injected group. Moreover, the development of embryo (E15) into neonatal stages (post 3 days), the expression of wdfy1 was increased oppositing PRDX6 expressed. These data indicate that wdfy1, at least play a significant role in the inhibition of neurogenesis in PRDX6 overexpressed mice brain.

TLR protein is present in mammalian brain cells in early embryonic stages of development and serves as a negative modulator of early embryonic neural precursor cell proliferation and axonal growth. Several members of TLR pathway including AKT1, CD14, Fos, IFNAR1, IL8, JUN, MAPK10, MAPK11, MAPK3, PIK3R1, PIK3R2, RELA and SPP1 are more highly expressed in descending neurons (DNs) (Fathi et al., 2011). Interestingly, it has been demonstrated that PRDXs is involved in TLR pathway, therefore, we investigated a link between PRDX6 and wdfy1 through TLR pathway for the control of neurogenesis in PRDX6 Tg mice. In the PRDX6 overexpressed mice brain, the expression of TLR4, TLR5, TLR6, TLR8 and TLR9 was slightly decreased whereas TLR3 and TLR7 was slightly increased. However, only TLR4 and TLR5 were significantly decreased. In further study, TLR4 expression was significantly increased in postnatal mice brain reversely consistent with PRDX6 expression. However, TLR5 was not changed in development stages. So, we focused on the TLR4 for further research. Moreover, knockdown of TLR4 significantly reduced neurogenesis, but PRDX6 expression was significantly increased with down regulation of PRDX6. Taken together, these study indicated that PRDX6 inhibits neurogenesis of neural stem cells through downregulation of wdfy1 mediated TLR4 signaling pathway, and suggest that the inhibitory effect of PRDX6 on neurogenesis may be at least play a role in the development of neurodegenerative disease.

## Innovation

Impaired neurogenesis has been associated with brain disorders. The role of peroxiredoxin 6 (PRDX6) in the neurodegenerative diseases is very controversial. To demonstrate the role of PRDX6 in neurogenesis, we compared neurogenesis ability of PRDX6 overexpressing transgenic (Tg) mice and wild type mice, and studied the molecular mechanisms. This study indicated that PRDX6 inhibits neurogenesis of neural precursor cells through TLR4 dependent downregulation of wdfy1.

## Materials and methods

### Animals and isolation of neural stem cells

The PRDX6 overexpressed tg mice were purchased from the Jackson Laboratory (Bar Harbor, ME, USA). The genetic background of non-transgenic mice and PRDX6 overexpressed mice is C57BL/6. The mice were housed and bred under specific pathogen-free conditions at the Laboratory Animal Research Center of Chungbuk National University, Korea (CBNUA-929-16-01). The mice were maintained in a room with a constant temperature of 22 ± 1 °C, relative humidity of 55 ± 10% and 12-h light/dark cycle and fed standard rodent chow (Samyang, Gapyeong, Korea) and purified tap water ad libitum. Neural stem cells were isolated from embryonic day 14 forebrain germinal zones from PRDX6 Tg or C57BL/6 (Jackson Laboratories). Bulk cultures were established and the medium included DMEM/F12, 10% FBS and 1% penicillin/streptomycin. After 24hr, the medium was changed with Neurobasal medium containing 1% glutamate, B27 supplement, 100 U/ml penicillin and 100 μg/ml streptomycin.

### PC12 cell culture and neurite outgrowth

PC12 cells were obtained from the American Type Culture Collection (Manassas, VA, USA). RPMI1640, penicillin, streptomycin, fetal bovine serum (FBS), horse serum (HS) and nerve growth factor (NGF) were purchased from Invitrogen (Carlsbad, CA, USA). PC12 cells were grown in RPMI1640 with 5% FBS, 10% HS, 100 U/ml penicillin and 100 μg/ml streptomycin at 37 °C in 5% CO2 humidified air.

### Neurite outgrowth assay

To study neurite outgrowth, the medium was changed to RPMI containing 1% HS, 100 ng/ml NGF, 100 U/ml penicillin and 100 μg/ml streptomycin. The cells were further cultured for 5 days. Cells with at least one neurite longer than two-body length were counted as neurite positive. At least 500 cells were counted for each group performed in triplicate.

### Differentiation of neural stem cells into motor neurons

For in vitro priming, neural stem cells were cultured in neurobasal medium plus B27 (Invitrogen), 0.1 mM 2-mercaptoethanol, 20 ng/ml b-fibroblast growth factor, 1 μg/ml laminin, 5 μg/ml heparin, 10 ng/ml neural growth factor (Invitrogen), 10 ng/ml sonic hedgehog (R&D Systems), 10 μM forskolin (Sigma) and 1 μM retinoic acid (Sigma) for 5 days. For the induction of astrocyte, neural stem cells were cultured in poly-L-lysine coated slides with DMEM medium plus N2 supplement and L-glutamate. All cultures were maintained in a humidified incubator at 37°C and 5% CO2 in air, and half of the growth medium was replenished every third day.

### Transfection

PC12 cells were transiently transfected with pcDNA or PRDX6 plasmid (Santa Cruz Biotechnology) using the WelFect-EX PLUS reagent in OPTI-MEN, according to the manufacturer’s specification (WelGENE, Seoul, Korea).

### Immunofluorescence staining

The fixed cells and tissues were permeabilized by exposure to 0.1% Triton X-100 for 2 min in PBS, and placed in blocking serum (5% bovine serum albumin in PBS) at room temperature for 2 h. The cells and tissues were then exposed to primary mouse monoclonal antibody for PRDX6 (1:200 dilution) and rabbit polyclonal antibody for Tubulin β-III (TUBBIII) (1:200) overnight at 4°C. After washes with ice-cold PBS, followed by treatment with an anti-mouse secondary antibody labeled with Alexa Fluor 488 and anti-rabbit secondary antibody labeled with Alexa Fluor 568 (1:100 dilution, Molecular Probes Inc., Eugene, OR) for 2 h at room temperature, immunofluorescence images were acquired using a confocal laser scanning microscope (TCS SP2, Leica Microsystems AG, Wetzlar, Germany) equipped with a 400× oil immersion objective.

### Western blot

The membranes were immunoblotted with primary specific antibodies. The blot was then incubated with corresponding conjugated anti-rabbit and anti-mouse immunoglobulin G-horseradish peroxidase (1:2000 dilution, Santa Cruz Biotechnology Inc.). Immunoreactive proteins were detected with the ECL western blotting detection system. The relative density of the protein bands was scanned by densitometry using MyImage (SLB, Seoul, Korea), and quantified by Labworks 4.0 software (UVP Inc., Upland, CA, USA).

### Immunoprecipitation

Neural stem cells were gently lysed for 1 hr on ice and then centrifuged at 14,000 rpm and 4 °C for 15 min, and the supernatant was collected. The soluble lysates were incubated with anti-PRDX6 antibody (Santa Cruz Biotechnology Inc) at 4 °C for o/n and then with Protein A/G bead (Santa Cruz Biotechnology Inc) for 4 h at 4°C and washed 3 times. Immune complexes were eluted by boiling for 5 min at 95 °C in SDS sample buffer followed by immunoblotting with anti-p21 (1:500) antibody.

### Immunohistochemistry staining

After being transferred to 30 % sucrose solutions, brains were cut into 20-μm sections by using a cryostat microtome (Leica CM 1850; Leica Microsystems, Seoul, Korea). After two washes in PBS (pH 7.4) for each 10 min, endogenous peroxidase activity was quenched by incubating the samples in 3 % hydrogen peroxide in PBS for 20 min, and two washes in PBS for each 10 min. The brain sections were blocked for 1 h in 5 % bovine serum albumin (BSA) solution and incubated overnight at 4 °C with a mouse polyclonal antibody against p21 (1:200; Santa Cruz Biotechnology, Inc., Santa Cruz, CA, USA), glial fibrillary acidic protein (GFAP) (1:200; Santa Cruz Biotechnology, Inc., Santa Cruz, CA, USA), Tubulin β-III (TUBBIII) (1:200; Cell Signaling Technology, Inc., Beverly, MA, USA) and a rabbit polyclonal antibody against Tyrosine hydroxylase (1:200; Cell Signaling Technology, Inc., Beverly, MA, USA). After incubation with the primary antibodies, brain sections were washed thrice in PBS for each 10 min. After washing, brain sections were incubated for 1 h at room temperature with the biotinylated goat anti-rabbit or goat anti-mouse IgG-horseradish peroxidase (HRP) secondary antibodies (1:500; Santa Cruz Biotechnology, Inc., Santa Cruz, CA, USA). Brain sections were washed thrice in PBS for each 10 min and visualized by chromogen DAB (Vector Laboratories) reaction for up to 10 min. Finally, brain sections were dehydrated in ethanol, cleared in xylene, mounted with Permount (Fisher Scientific, Hampton, NH), and evaluated on a light microscopy (Olympus, Tokyo, Japan) (×50 or ×200).

### Immunofluorescence staining

For Immunofluorescence staining, the fixed cells and brain sections were exposed to the following primary antibodies: Tubulin β-III (TUBBIII) (1:200; Cell Signaling Technology, Inc., Beverly, MA, USA) and Neurofilament (1:200; Cell Signaling Technology, Inc., Beverly, MA, USA) at room temperature for 2 h. After incubation, the cells were washed twice with ice-cold PBS and incubated with an anti-mouse or mouse secondary antibody conjugated to Alexa Fluor 488 or 568 (Invitrogen-Molecular Probes, Carlsbad, CA) at room temperature for 1 h. Immunofluorescence images were acquired using an inverted fluorescent microscope Zeiss Axiovert 200 M (Carl Zeiss, Thornwood, NY) (×200).

### Neural stem cell SVZ injections

Wdfy1 siRNA transfected neural stem cells or EV control (30,000 particles in 1 mL, four animals per condition) were stereotaxically injected into the dorsal horn of the SVZ of 10-wk-old ICR mice using coordinates AP +0.5; ML +1.1; from bregma and DV -1.9 from the pial surface. Mice were sacrificed 2 wk following injection, and the CNS tissue was fixed by transcardial perfusion with PBS followed by 4% paraformaldehyde. To count cells in the olfactory bulb, five different fields at 203 were used from each animal.

### RNA preparation

Total RNA was extracted from the mouse neural stem cells using the TRI REAGENT (MRC, OH) according to the manufacturer’s instructions. Following homogenization, 1 ml of solution was transferred to a 1.5 ml Eppendorf tube and centrifuged at 12,000 g for 10 minutes at 4°C to remove insoluble material. The supernatant containing RNA was collected, mixed with 0.2 ml of chloroform, and centrifuged at 12,000 g for 15 minutes at 4°C. After RNA in the aqueous phase was transferred into a new tube, the RNA was precipitated by mixing 0.5 ml of isopropyl alcohol and recovered by centrifuging the tube at 12,000 g for 10 minutes at 4°C. The RNA pellet was washed briefly in 1 ml of 75% ethanol and centrifuged at 7,500 g for 5 minutes at 4°C. Finally, the total RNA pellet was dissolved in Nuclease-water, and its quality and quantity was assessed by Agilent bioanalyzer 2100 analysis. Gene expression was analyzed with GeneChip^®^ Human Gene 2.0 ST Arrays (Affymetrix, Santa Clara, CA), which is comprised of over 21,000 protein coding transcripts and over 19,000 entrez genes. For each gene, eleven pairs of oligonucleotide probes are synthesized in situ on the arrays.

### Microarray

Fragmented and Labeled ss-DNA were prepared according to the standard Affymetrix protocol from 400ng total RNA (GeneChip^®^ WT PLUS Reagent Kit Manual, 2001, Affymetrix). Following fragmentation, 3.5 ug of ss-DNA were hybridized for 16 hr at 45C and 60 rpm on GeneChip^®^ CHO Gene 2.0 ST Array. GeneChips were washed and stained in the Affymetrix Fluidics Station 450. GeneChips were scanned using the Affymetrix GeneChip Scanner 3000 7G. The data were analyzed with Robust Multichip Analysis (RMA) using Affymetrix default analysis settings and global scaling as normalization method. The trimmed mean target intensity of each array was arbitrarily set to 100. The normalized, and log transformed intensity values were then analyzed using GeneSpring GX 13.1 (Agilent technologies, CA). Fold change filters included the requirement that the genes be present in at least 200% of controls for up-regulated genes and lower than 50% of controls for down-regulated genes. Hierarchical clustering data were clustered groups that behave similarly across experiments using GeneSpring GX 13.1 (Agilent technologies, CA). Clustering algorithm was Euclidean distance, average linkage.

### Quantitative real-time PCR (qPCR)

For mRNA quantification, total RNA was extracted using the easy-BLURTM total RNA extraction kit (iNtRON Biotech, Daejeon, Korea). cDNA was synthesized using High Capacity cDNA Reverse Transcription Kits (Applied Biosystems, Foster city, CA) according to the manufacturer’s instructions. Briefly, 2 μg of total RNA was used for cDNA preparation. Quantitative real-time PCR was performed using the Brilliatn III Ultra-Fast Green QPCR Master Mix (Agilent Technologies, Waldbronn, Germany) specific for 18S and wdfy1 (5’-ACCATCCGAGTATGGCTGAAA-3’ and 5’-CCTGCTGTCGTGGTGGTATG-3’). All reverse transcription reactions were run in a StepOnePlus Real-Time PCR System (Applied Biosystems, Foster city, CA) using the universal cycling parameters (3 min 95 °C, 40 cycles of 5 s 95 °C, 12 s 60 °C). Results were normalized to 18S and quantified relative to expression in control samples. For relative quantification calculation, the 2^-ΔΔ CT^ formula was used, where: -ΔΔ CT = (C_T,target_ - C_T,18S_) experimental sample - (C_T,target_ - C_T,18S_) control sample.

### Statistical analysis

The data was analyzed using the GraphPad Prism 4 ver. 4.03 software (GraphPad Software, La Jolla, CA). Data is presented as mean ± SD. The homogeneity of variances was assessed using the Bartlett test. The differences in all data were evaluated by one-way analysis of variance (ANOVA). When the *P* value in the ANOVA test indicated statistical significance, the differences were evaluated by the Dunnett’s test. The value of *P* < 0.05 was considered to be statistically significant.

## Acknowledgement

This work was supported by the National Research Foundation of Korea (NRF) grant funded by the Korea government (MSIP) (No. MRC, 2008-0062275).

## Author contributions

MHP and JTH contributed to the design and coordination of the study. MHP performed all experiments. MHP, DJS and KTN participated in the study design and prepared the manuscript. SYK, SYO and MJS helped with image analysis and microscopy. HKC and THL participated in the technical supports. All authors read and approved the final manuscript prior to submission.

